# CRISPR-Cas12b-assisted nucleic acid detection platform

**DOI:** 10.1101/362889

**Authors:** Linxian Li, Shiyuan Li, Jin Wang

## Abstract

Rapid molecular diagnostic technology is very useful in many areas, including public health, environmental testing and criminal investigation. We recently showed that Cas12a had *trans*-cleavage activity upon collateral single-stranded DNA (ssDNA), with which the HOLMES platform (one-<u>HO</u>ur <u>L</u>ow-cost <u>M</u>ultipurpose highly <u>E</u>fficient <u>S</u>ystem) was developed. Here, we combine the thermophilic Cas12b, which also has the ssDNA *trans*-cleavage activity, with Loop-Mediated Isothermal Amplification (LAMP), and create HOLMESv2. In HOLMESv2, LAMP amplification and Cas12b *trans*-cleavage can be integrated into a one-step system with a constant temperature, which therefore brings much convenience in nucleic acid detection. Moreover, we also simplify the RNA detection procedures in HOLMESv2, using an RNA-dependent DNA polymerase for amplification and therefore omitting an extra reverse transcription step.

**One Sentence Summary:** We combine LAMP and Cas12b to develop HOLMESv2 for conveniently detecting target nucleic acid in a one-step approach.

## Main Text

Due to the high sensitivity, specificity and simplicity, the recently emerged CRISPR-based nucleic acid detection methods have shown great promise in clinical diagnostics and have therefore drawn much attention (*1, 2*). The principle of these methods is based on the *trans*-cleavage activities of some Cas proteins, including the DNA-targeting and single-stranded DNA (ssDNA)-cleaving Cas12a (*3-8*). Combined with amplification, Cas12a has been employed to develop HOLMES (one-HOur Low-cost Multipurpose highly Efficient System) or DETECTR (DNA Endonuclease Targeted CRISPR *Trans* Reporter) (*3, 6, 7*). However, in both systems, the amplification and detection processes are separated, which might cause practical inconvenience. Here, we employ Cas12b (previously known as C2c1) (*9, 10*), a thermophilic RNA-guided endonuclease from type V-B CRISPR-Cas system, to develop an integrated one-step method in one-pot reaction system, termed HOLMES version 2 (HOLMESv2).

We first purified the *Alicyclobacillus acidoterrestris* Cas12b protein (AacCas12b) (*9*), and verified that Cas12b had cis-cleavage activity against targeted double-stranded DNA (dsDNA) (**Fig. S1**). We then tested the Cas12b *trans*-cleavage activity upon a 12-nt ssDNA probe, which was labeled with fluorescence HEX on the 5’ end and quencher BHQ1 on the 3’ end (HEX-N12-BHQ1), and confirmed that both double-stranded DNA (dsDNA) and ssDNA targets were able to trigger Cas12b to *trans*-cleave the ssDNA probe (**Fig. 1a and 1b**), which was consistent with the previous findings both by our group (*11*) and by Chen *et al*. (*3*). Besides, we found that the Cas12b ternary complex with ssDNA showed higher *trans*-cleavage activity and the fluorescence reached the highest value in about 10 min, while at least 30 min was required with dsDNA (**Fig. 1c**). Notably, the target preference of Cas12b was distinct from that of Cas12a, which showed higher *trans*-ssDNA cleavage rate with dsDNA target (*3*).

**Fig. 1.**
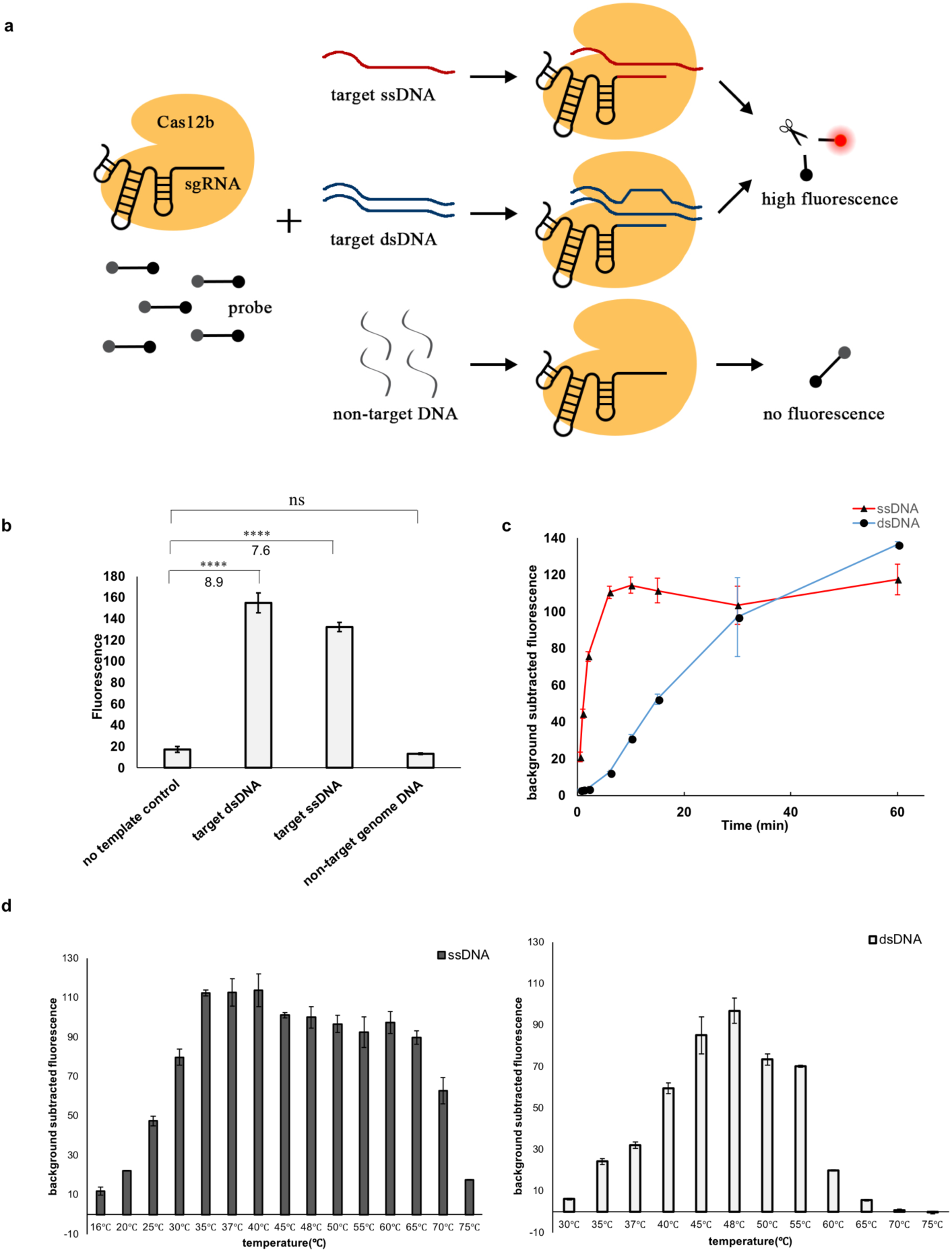
Cas12b has *trans*-cleavage activity on collateral ssDNAs. (**a**) Schematic of Cas12b *trans*-cleavage activity. (b) The fluorescence signal was detected after incubating Cas 12b, sgRNA (sgRNA-T1), target DNA and collateral ssDNA probe (HEX-N12-BHQ1) at 48 °C for 30 min. (c) Time-course (0-60 min) analysis of the Cas12b *trans*-cleavage activity with target ssDNA or target dsDNA. (d) Analysis of the temperature range required for Cas12b *trans*-cleavage with either target ssDNA or target dsDNA. Error bars represented the mean ± s.d. (n = 3 replicates).

According to the previous study (*10*), the PAM site of “5’-TTN-3’” is required by AacCas12b to cleave target dsDNA. To test whether PAM is required for *trans*-cleavage by Cas12b, we designed different sequences (e.g. 5’-TTC-3’, 5’-TAC-3’, 5’-ATC-3’, 5’-AAC-3’, 5’-GGC-3’ and 5’-CCC-3’) at the PAM position (**Fig. S2a**). As shown in **Figure S2b**, with target dsDNA, the Cas12b-mediated *trans*-cleavage was sensitive to the PAM sequence, and only with the PAM sequences of 5’-TTC-3’ and 5’-TAC-3’ Cas12b showed high *trans*-cleavage activity, which was reflected by the high fluorescence signal. However, when Cas12b targeted ssDNA substrates, it *trans*-cleaved the ssDNA probe in a PAM-independent manner (**Fig. S2b**). Then, we tested different temperatures required for the Cas12b *trans*-cleavage reaction, and found the suitable temperature ranged from 35 °C to 65 °C with target ssDNA and from 45 °C to 55 °C with target dsDNA (**Fig. 1d**).

Using the above fluorescence reporter system, we next determined the lowest target concentration required for triggering the Cas12b *trans*-cleavage, which was as low as 1 nM for either target ssDNA or target dsDNA (**Fig. S3**). However, when combined with isothermal DNA amplification method (e.g. Loop-Mediated Isothermal Amplification (LAMP) (*12*)), the target concentration could decrease to 10^−8^ nM (**Fig. 2a**), which was the same as that of Cas12a (*7*). Therefore, because of the high sensitivity for DNA substrate, Cas12b can also be employed for nucleic acid detection, and the new method combining both amplification and Cas12b *trans*-cleavage detection is named HOLMESv2.

**Fig. 2.**
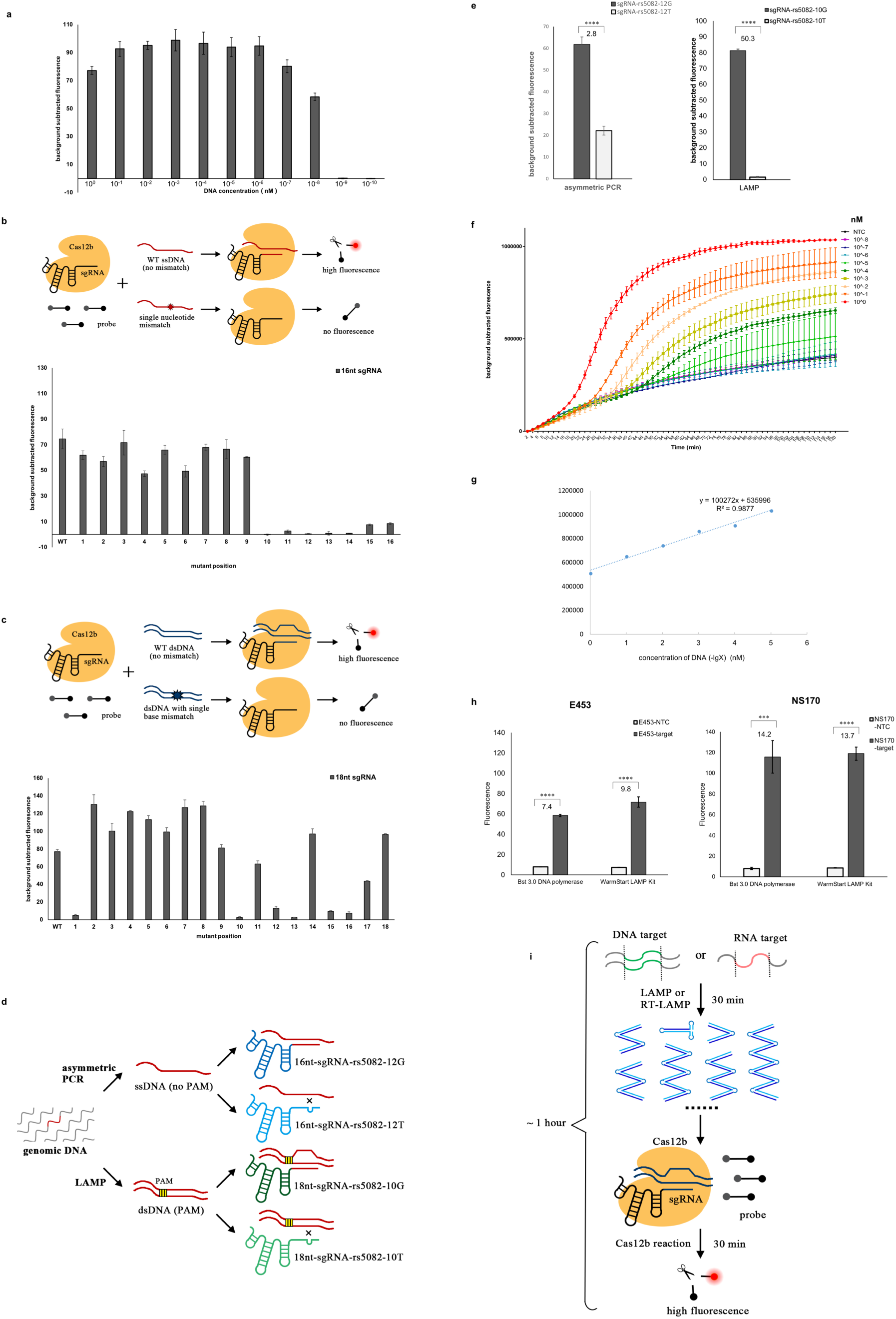
Target nucleic acid detection by HOLMESv2 (combined nucleic acid amplification and Cas12b detection). (**a**) Determination of HOLMESv2 sensitivity. Target DNA (pUC18-DNMT1-3) was serially diluted, amplified by LAMP amplification and followed by Cas12b *trans*cleavage, and ffluorescence was determined with a fluorescence reader. (**b**) Distinguishing ssDNAs with single-nucleotide mismatch by Cas12b. The schematic was illustrated (upper panel), and the results (bottom panel) were obtained with the use of an sgRNA with 16-nt wildtype guide sequence. Mutated target sequences were listed in **Figure S4a**, and only mutated sequences of 1 to 16 were analyzed with the 16-nt sgRNA. (**c**) Distinguishing dsDNAs with single-base mismatch by Cas12b. The schematic was illustrated (upper panel), and the results (bottom panel) were obtained with the use of an sgRNA with 18-nt wild-type guide sequence. Mutated target sequences were listed in **Figure S5a**, and only mutated sequences of 1 to 18 were analyzed with the 18-nt sgRNA. (**d**) Schematic of SNP distinguishing by HOLMESv2. Target DNA containing the SNP locus (e.g. rs5082) was either amplified by asymmetric PCR or LAMP. For LAMP amplification, a PAM sequence was required, which could be introduced by primers if there was no suitable PAM nearby. (**e**) Genotyping the rs5082 locus in the human cell HEK293T by HOLMESv2. The target region was either amplified by asymmetric PCR (left panel) or LAMP (right panel). For asymmetric PCR, the sgRNA was designed to align the mismatch at the 12^th^ position of the guide sequence, counting from the PAM sequence. While for LAMP, the mismatch was aligned at the 10^th^ position. (**f**) Quantitation of target DNA by one-step HOLMESv2. Serially diluted plasmid pUC18-rs5082 was used as the targets, and FAM-N12-Eclipse was used as the probe with FAM fluorescence real-time detected by a real-time machine. (**g**) Standard curve analysis of the one-step HOLMESv2. Fluorescence values were collected for targets ranging from 1 nM to 10^−5^ nM at the time point of 120 min. (**h**) Detection of RNA virus by HOLMESv2. Japanese encephalitis virus (JEV) RNA was amplified by either the reverse transcription LAMP (WarmStart LAMP Kit) or merely Bst 3.0 DNA polymerase, followed by Cas12b detection. Reactions with no template added were designated as NTC. Two different sgRNAs were designed, targeting E453 and NS170 sites in JEV, respectively. (**i**) Schematic of nucleic acid detection by HOLMESv2 with both LAMP (or RT-LAMP) and Cas12b detection.

We then wondered whether HOLMESv2 could distinguish target DNAs with single-nucleotide mismatch. Considering the ability to distinguish single-nucleotide mismatch by Cas12a is determined by with both the length of the CRISPR RNA (crRNA) guide sequence and the position of the mismatch (*7*), we analyzed both sgRNAs with distinct guide length and target ssDNA (or dsDNA) sequences with single nucleotide polymorphisms (SNPs) at different positions. We first tested target ssDNA (DNMT1-3), and found only sgRNA with 16-nt or 17-nt guide sequences (abbreviated as 16-nt or 17-nt sgRNA thereafter) showed great differences in fluorescence signals when the mismatch was aligned within the 10^th^ - 16^th^ positions (**Fig. 2b** and **Fig. S4a, S4b**). Otherwise, Cas12b either failed to distinguish targets with SNP (e.g. all with high signals when using 18-nt - 20-nt sgRNAs) or produced remarkably reduced *trans*-cleavage activity (e.g. using 14-nt - 15-nt sgRNAs) with all targets (**Fig. S4b**). In addition, we also tested the 16-nt sgRNA for other ssDNA targets and demonstrated it effective for discriminating SNPs (**Fig. S4c** and **S4d**); however, sgRNAs should be designed to align the mismatch at different positions of the guide sequences for distinct targets (e.g. 8^th^ and 10^th^ - 12^th^ for rs5082, 11^th^ for rs1467558 and 8^th^ - 15^th^ for rs2952768) (**Fig. S4d**). We then tested target dsDNA of DNMT1-3 (**Fig. S5a**), and found 18-nt - 20-nt sgRNAs were more effective for Cas12b to discriminate single-base mismatch (**Fig. 3c** and **Fig. S5b**). However, different guide length led to biased preferences for the positions of the mismatch. For example, with the 18-nt sgRNA, Cas12b was able to distinguish the dsDNA target of DNMT1-3 with single-base mismatch positioned at 1^st^, 10^th^, 12^th^, 13^th^, 15^th^ and 16^th^ of the guide sequences, but positions changed when either 19-nt or 20-nt sgRNAs was used (**Fig. 3c** and **Fig. S5b**). Therefore, HOLMESv2 is of both high sensitivity and high specificity, where the sensitivity relies on the amplification methods and the specificity is determined by Cas12b.

Based on the above findings, we next used HOLMESv2 to examine the SNP locus of rs5082 from human cell HEK293T. As there is no suitable PAM sequence nearby the rs5082 locus, we employed two distinct approaches to amplify the target, including asymmetric PCR and LAMP. When combined with asymmetric PCR, which generates ssDNA products, Cas12b could distinguish the SNP locus without a PAM sequence, and only with perfectly matched sgRNA

Cas12b produced high *trans*-cleavage activity (**Fig. 2d and 2e**). But with LAMP amplification, which produces dsDNA products, the PAM sequence is required and was therefore introduced by primers (**Table S3**) to assist Cas12b in distinguishing the SNP locus (**Fig. 2d** and **2e**).

Moreover, we tested the practicability of integrating two steps of isothermal LAMP amplification and Cas12b detection into one reaction system, where the amplification and detection performed at the same time. Because both steps have a wide temperature range, the integrated system was tested under temperature ranging from 48 °C to 60 °C to find out the most suitable temperature for the one-step HOLMESv2. As shown in **Figure S6**, with the increase of the reaction temperature, both the period of lag phase and the fluorescence signal that reflected the Cas12b *trans*-cleavage activity reduced, and the signal was hardly detectable when the temperature was above 57.8 °C, which was consistent with the temperature range suitable for Cas12b *trans*-cleavage (**Fig. 1d**). Therefore, temperature of 55 °C was finally selected for HOLMESv2 one-step system. Assisted with a real-time fluorescence reader, we could real-time monitored the one-step HOLMESv2 reaction and quantitated target DNA (i.e. plasmid pUC18-rs5082), which was serially diluted to concentrations ranging from 1 nM to 10^−8^ nM. As expected, higher target concentration would lead to less take-off time and higher fluorescence signal (**Fig. 2f**), and with the fluorescence values at the time point of 120 min, a standard curve was generated with the R^2^ value of 0.9877 (**Fig. 2g**). Although we here proved the practicability of one-step HOLMESv2, the lowest target concentration within the linear range was 10^−5^ nM under this tested condition, which was much less than the sensitivity of the separated HOLMESv2 system. Therefore, further improvements might be needed, which include both refinement of the reaction buffer and chemical modification of the enzymes to reduce cross interference.

Besides of DNA, RNA can also be detected with the CRISPR-based methods; however, RNA needs to be first reverse transcribed into cDNA, which is then used as the template for either *in vitro* transcription (e.g. SHERLOCK or SHERLOCKv2) or amplification (HOLMES or DETECTR) (*3-8*). Although reverse transcription and subsequent amplification can be integrated into a one-pot system, the whole process contains more than one enzyme and can be complicated. We therefore used DNA polymerases that contained the 5’ → 3’ DNA polymerase activity with either DNA or RNA templates (e.g. the engineered Bst DNA polymerase (*13*)) to directly amplify the RNA target, which could simplify the RNA detection procedure by omitting an extra reverse transcription step. Similar to the reverse transcription LAMP (RT-LAMP) reaction system, HOLMESv2 with LAMP using merely Bst 3.0 DNA polymerase successfully detected the Japanese encephalitis virus (JEV) within one hour (**Fig. 2h**). Therefore, using fewer enzymes, HOLMESv2 can be more robust and convenient for detection of RNA targets (**Fig. 2i**).

In theory, Cas12b detection can be coupled with any amplification methods to improve both the sensitivity and specificity. Considering many isothermal amplification methods (e.g. LAMP (*12*), RPA (*14*), NASBA (*15*) and CPA (*16*)) have been successfully tested for amplification of single nucleic acid molecule, HOLMESv2 therefore has great potential for further clinical diagnostics at the single-molecule level. In addition, HOLMESv2 can also be integrated into present diagnostic instruments and platforms, including the portable lateral flow assay devices for convenient detection (*17*), the microfluidic platform for multiple targets (*18, 19*), the quantitative PCR machine for relative quantification and the digital PCR system for absolute quantification (*20*).

## Acknowledgments

We thank Guanghui Xie for help in analyzing the real-time HOLMESv2 data, Dawei Chen for help in expanding the applications of HOLMESv2, Xuan Zheng for help in drawing illustration charts, and members of Tolo Biotech. Co., Ltd. for comments and discussions.

## Author contributions

L.L. and S.L. contributed equally to the work and conducted the majority of the experiments. J.W. conceived the study and S.L. designed the experiments. S.L. and J.W. wrote the manuscript.

## Data and materials availability

All data are available in the manuscript or the supplementary material.

## Supplementary Materials

Materials and Methods

Figures S1-S6

Tables S1-S3

